# Study on H3K9 acetylation modification and TLR9 immune regulation mechanism in patients with anti-HBV treatment using thymosin a1 combined with entecavir

**DOI:** 10.1101/2020.10.31.363127

**Authors:** Hai-Peng Zhu, Ke Wang, Wei Du, Huan-Huan Cao, Qing-Yang Zhong, Si-Chun Yin, Jian-Bo Zhong, Fa-Wu Li

## Abstract

For hepatitis B antiviral treatment, there has been no comprehensive method yet. Interferon has poor antiviral efficacy, while nucleoside drugs have long course of treatment and high relapse rate. To improve the anti-HBV curative effect, treatment methods such as thymosin combined with entecavir have become a focus of clinical investigation. To explore potential mechanism of the combination therapy, based on previous studies, this paper explores the relationship between TLR9 expression in PBMCs, secretion of corresponding downstream inflammatory factors and HBV load in anti-HBV treatment with Thymosin a1 (Ta1) combined with entecavir. Chromatin immunoprecipitation combined with PCR method was adopted to detect H3K9 acetylation modification in patients. The relationship between TLR9 expression was explored using RT-QPCR, the relationship between secretion of inflammatory factors, efficacy and TLR9 mRNA expression was determined using Luminex technology. The results showed that during anti-HBV treatment with Ta1 combined with entecavir, histone acetylation increased in patients’ PBMCs, acetylated protein H3K9Ac had significant binding with promoter region of the TRL9 gene, thereby increasing the expression of TRL9 mRNA, activating the immune pathway under TRL9 regulation, promoting secretion of inflammation factors IL-6, IL-12, IFN-γ, and TNF-α, boosting the progress of antiviral therapy. H3K9 acetylation modification of TLR9 exists and plays an important role in patients with chronic hepatitis B. During the combination therapy with entecavir and Ta1, histone acetylation modification of TLR9 was significantly improved, which increased the expression of TLR9 at the mRNA and protein levels, and further regulated IL-6, IL-12 and other cytokines.

## 1. Introduction

China is a big country of liver disease (1-3). Statistics show that there are nearly 100 million people with chronic hepatitis B virus (HBV) infection in the country (4). Where, HBV infection rate is particularly high in Guangdong (5). About 20% patients may deteriorate to cirrhosis, liver cancer or liver failure. Chronic hepatitis B has complex pathogenesis (2). Current treatments mainly include antiviral therapy, immune regulation, anti-inflammation and oxidation, and anti-fibrosis. Where, entecavir as a potent and low-resistance anti-HBV treatment has become a recognized approach (6), but it has the shortcomings of limited efficacy, long-term treatment and high relapse rate after drug withdrawal (7). Seen from clinical perspective, it is necessary to further improve its efficacy and shorten the course of treatment. A number of anti-HBV studies on Thymosin a1 (Ta1) combined with entecavir have indicated that this treatment regimen can improve serum ALT recovery rate, HBV DNA negative conversion rate, serum HBsAg negative conversion rate, and serum HBeAg/anti-HBe conversion rate (8-11). Moreover, there are few and slight clinical side effects (12). Entecavir exerts direct antiviral effect mainly by inhibiting the activation of HBV polymerase, inhibiting the formation of negative strand by reverse transcription of pre-genomic mRNA, and inhibiting DNA-dependent DNA synthesis (13). Ta1 mainly regulates the body’s immune function for antivirus (14-15). Therefore, it is very important to study antiviral mechanism and response model of chronic hepatitis B with treatment of chronic hepatitis B using thymosin combined with entecavir as the research carrier.

Toll-like receptors (TLRs) are an important class of receptors that recognize foreign microorganisms in innate immunity, which can initiate direct killing effect of innate immunity and mediate secondary onset of adaptive immunity (16-17). TLR9 is one of the receptors with multiple effects and big influence (18). Ta1 can improve the body’s anti-infective ability by activating TLR signaling pathway. Studies have found that Ta1 can up-regulate TLR9 expression by plasma cell-like DCs, thereby inducing the expression of IL-10 and IL-12. It can also obtain anti-fungal innate immunity and protective Th1 cell immune response by activating signaling pathways such as TLR2 and TLR9 (19). Moreover, it can also greatly enhance pDC ability to secrete INF-α through TLR9 signaling pathway, thereby clearing CMV (20). However, in the anti-HBV infection treatment, there is no relevant research on whether Ta1 affects TLR9 expression.

Earlier, when studying the role of TLR1-10 in chronic hepatitis B, it was found that only TLR9 and TLR10 expressions were related to HBV load, suggesting that these two recognition receptors are closely related to HBV replication (21-22). TLR9 extracellular region is a key region for identifying foreign microorganisms (23). Capable of identifying non-methylated CpG gene sequences, it has the ability to recognize HBV virus and initiate immune response. Recent studies have shown that TLR9 may be involved in the development of chronic hepatitis B disease (24). Xu et al. found that compared with normal people, patients with chronic hepatitis B had lower TLR9 mRNA expression in peripheral blood mononuclear cells (25). It is considered that HBV might inhibit TLR9 mRNA expression in chronic hepatitis B patients’ PBMc as a certain starter factor, eventually causing immune escape or immune tolerance. This view was also confirmed by Vincent et al (26). who found in HBV-infected people’s pDC that HBV could down-regulate the expression of TLR9 mRNA and protein to reduce the production of INF-α, which in turn caused persistent virus infection. Moreover, for TLR9, in addition to the traditional regulation, there are also epigenetic modifications. Numerous reports show that acetylation modification of histone H3 and lysine 9 (H3K9) is related to gene transcription development, while deacetylation is related to gene inactivation (27-29). Importantly, it has been reported in the literature that histone acetylation modification state is closely related to various viral infections. Earlier, the group compared acetylation modification state of histone H3K9 of peripheral blood CD4^+^T cells in whole genome promoter region in different disease states of chronic hepatitis B, finding that H3K9 acetylation modification regulation exists in two sequence regions of TLR9, suggesting that H3K9 acetylation modification of TLR9 may also play an important role in the occurrence and development of chronic hepatitis B (21).

At present, for the treatment of chronic hepatitis B based on thymosin and entecavir, exploration into whether TLR9 affects the body’s immunity, whether HBV virus regulates TLR9 expression, whether TLR9 is related to the therapeutic effect, and the mechanism of TLR9 in hepatitis B patients will provide an important theoretical basis for determining the pathogenesis of chronic hepatitis B and clinical antiviral mechanism. To continue our research in the field of chronic hepatitis B (21, 30-31), using real-time PCR, Luminex, and Chip technology, this paper explores the relationship between TLR9 expression in PBMCs, corresponding downstream inflammatory factor secretion levels and HBV load in anti-HBV treatment with Ta1 combined with entecavir from two aspects: clinical treatment and immune mechanism. It is confirmed that TLR9 is involved in the immune response in anti-HBV process using Ta1 combined with entecavir. The research results provide a new basis for the antiviral immune control theory of chronic hepatitis B, which has broad social needs and important clinical application significance.

## 2. Materials and methods

### 2.1 Research objects and sample collection

For patients with chronic hepatitis B (serum HBsAg positive for more than 6 months) (18-65 years old) who were hospitalized in the Department of Infectious Diseases, Dongguan People’s Hospital from December 2017 to February 2019, their general and clinical data were collected and informed consent was signed. Inclusion criteria: The diagnosis complies with “Guidelines for the Prevention and Treatment of Chronic Hepatitis B” jointly developed by Hepatology Branch of the Chinese Medical Association and Infectious Disease Branch of the Chinese Medical Association in 2015. It also refers to “Prevention and Treatment Program for Viral Hepatitis” jointly revised by Infectious Diseases and Parasitic Diseases Branch, Hepatology Brach of Chinese Medical Association in 2000. The study chose those with serum HBsAg-positive, HBeAg-positive and HBV DNA>2×10E4 IU/mL or HBeAg-negative and HBV DNA>2×10E3 IU/mL; with no decompensated liver cirrhosis, no use of other antiviral drugs and immunizations within six months.

Exclusion criteria: The study excluded those with HAV, HCV, HDV, HEV, HIV and other viral infections; with HCC or AFP≥400ng/ml for more than 1 month; those need immunosuppressive treatment or radiotherapy, chemotherapy due to other diseases; those with positive pregnancy test or breastfeeding patients; patients who could not follow the study schedule and sign informed consent. Midway withdrawal criteria: Patients can withdraw from the study at any time before sample collection. The inclusion, exclusion and withdrawal criteria for the healthy control group: there should be no HBV infection, other exclusion and withdrawal criteria is the same as above.

This study has been reviewed by the Ethics Committee of Dongguan People’s Hospital (NO. 2017079). 5 mL venous blood was collected from the cubital vein and placed in an anticoagulant tube containing 500μL heparin. PBMCs anticoagulant tube was separated within 2 h after blood collection to collect blood samples from the subjects for TLR9 mRNA expression detection. TLR9 gene received H3K9 acetylation detection. At the same time, 5 mL blood sample was collected from the subject with an anticoagulation tube. The plasma was collected by centrifugation and stored at −20°C for the detection of inflammatory factors.

### 2.2 Treatment Information

Therapeutic drugs, Entecavir, Baraclude, Sino-American Shanghai Squibb Pharmaceutical Co., Ltd.; Thymosin a1, 1.6 mg injection, Thymalfasin, Chengdu Diao Pharmaceutical Group. All the samples were divided into 3 groups: 15 healthy volunteers (Control), 28 patients with chronic hepatitis B (CHB), and 29 cases treated with thymosin a1 combined with entecavir (Ta1-ETV) for three month. The three groups were comparable in terms of gender and age (P> 0.05). HBV load and HBsAg/HBeAg quantitative detection, HBV load was detected using HBV DNA quantitative detection kit of Daan Gene Co., Ltd., HBsAg/HBeAg quantitative detection was performed using Abbott’s chemical fluorescence detection kit.

### 2.3 TLR9 mRNA expression during combination therapy

PBMCs isolated from blood samples were extracted with total RNA using the Trizol method (total RNA Extraction Kit, DP431, Tiangen), and the purity NanoDrop2000, Thermo) was measured by an ultraviolet spectrophotometer. Follow the reverse transcription box instructions for reverse transcription (FastKing One Step RT-PCR Kit, KR123, Tiangen). Real-time PCR (Talent qPCR PreMix (SYBR Green), FP209, Tiangen) was performed using the designed TLR9 gene-specific primers. The primers and amplification conditions were: F-AACTGGCTGTTCCTGAAGTC, R-TGCCGTCCATGAATAGGA AG, annealing temperature 55°C, product length 394 bp; internal reference gene β-actin: F-AGGCCAACCGCG AAGATGACC, R-GAAGTCCAGGGCGACGTAGCAC, annealing temperature 55°C, product length 350 bp. The Ct data was obtained from the Bio-Rad PRISM Sequence Detection software in the Fluorescence quantitative PCR instrument.

### 2.4 Serum inflammatory factor detection by Luminex technology

Isolate and preserve serum samples, use multi-factor detection antibody chip platform, and adopt R & D Systems High Sensitivity (HS) Cytokine Premixed Magnetic Luminex Performance Assay kit (FCSTM09-04) to complete detection of four protein indicators IL-6, IL-12p70, IFN-γ, TNF-α in human serum samples. The experimental procedure is summarized as follows. The sample was diluted 5 times with the same dilution as the dilution standard substance, that is, 20 μL sample stock solution was added to the 80 μL diluent.

Prepare all required reagents and standard substances as described in the reagent preparation in the kit instructions. Remove the black transparent-bottomed microplate from the dense kit equilibrated to room temperature. Unused strips were sealed with a sealing plate film. Add standard substances and experimental samples of different concentrations to the corresponding wells, 50 μL each; resuspend and mix the prepared microsphere working solution, and add 50μL to each well. After the reaction wells were sealed with special sealing tape, place the solution in a microplate shaker, and incubate it at 800 rpm for 2 h with shaking at room temperature. Place the reaction plate in a hand-held magnetic stand, hold it for 1 min, and then shake off the liquid in the plate. Add 100 μL washing solution to each well, hold it for 1 min, and then shake it off. There is no need to pat dry it on absorbent paper. Repeat the washing operation twice and wash the plate 3 times.

Add 50 μL of biotinylated detection antibody working solution to each well, seal the reaction wells with a sealing plate, place it in a microplate shaker, and incubate it at 800 rpm for 1 h at room temperature with shaking. Add 50 μL of Strepavidin-PE working solution to each well, seal the reaction wells with a sealing plate, place it in a microplate shaker, incubate it at 800 rpm for 30 min with shaking at room temperature. Add 100 μL washing solution to each well to resuspend the microspheres, and place it on a microplate shaker, incubate at 800 rpm for 2 min with shaking at room temperature. Test it on a multi-factor analyzer (Luminex 200) within 90 min. According to the obtained fluorescence signal value and the concentration of each indicator substance, software Xponent3.1 was used to perform a five-parameter equation fitting and calculate the concentration of the unknown sample.

### 2.5 Chromatin immunoprecipitation and H3K9 acetylation detection of TLR9 gene

PBMCs in blood were separated from all fresh samples using PBMc separation solution, and the separation volume of each sample should be not less than 10 million cells. The predicted binding sites of TLR9 gene and H3K9 acetylation are as follows. >chr3:52226200-52226513 (reverse complement)

AGCCCCTGGAGGGCAGAGAGCAGGGCAGGACAGCCAGAAGGGAGGTGGCACCTGCATGGGTCTAGGAGAGAGGATGGGGGCTGGTGGACATGGGGACGGTGGGCTGTGGGCACACCCTTCAGCATGGTGGACCCAGCAGAACTTGCTGAGGGGCCCGGGGTCACCCAAGGCTTTGGGCCCGAGGAATCTGAGTCTCCTCACCTAGATCAGAAGGAGAGTGGGAAGAACTGATGGGGAGGACTGGCTGGCGGCCCTGCTGGGCCAGCACACACCTGGCCTCTAGGAGCCCCATCTGGAGTGACGTGGTGTGTGCT

>chr3:52226753-52227102 (reverse complement)

TGTACATAATTCAGCAGATATCAAGCACTTACTATGTGCTGGGCACTGTACTGGATCCTGGGGATGCAGATAAAAGATCACTGCCCTTAAGAAGCTGACATTC CAGCAGGGGAATAAGACGATATACAATAAACCATGAAAGATCAATGATCCGGTGTGCTAGCAGTTAAAAAATGTTAGGACAAAGAGAAACATAGACCAGGCAAAGGAGCTCAGGAGTGCCAGATCTGGGGTGGGAGGTTTGTAAGAAGGCTGGATGGCCCTGTTGAGAGGGTGACATGGGAGCAGAGACATAATGGAGGCAAAGGAGGGGTCATATGAGACTTGGGGGAGTTTTCAGGCAGAGGGAA

>chr3:52227754-52228023 (reverse complement)

TTCACCCTCCCTGCCCCCACCACCTACCCCTGTCAAGATGGGTAACTTAATCACATCTGCGAAGTCGTTTTTGCCACACTGTGGGGTGTTGGGGTCACGTGTTTGCAGGTTTGGGGAATTAGGACAAGGATCTCTGAGAGGGACTTTATGCAGCCTCCCACATGGGATAAGGGCTCCTCCTCGAAGGCTTCCCAGCCTCCCTGGGCTGAGGCCAGGACAAGTTTTTCTGTGGACATCGATATCGGTGTCTCCAAGCTGAGTGTGTCCATG

Add 100 μL cell lysis buffer to the PBMc cells isolated from the blood, followed by vortex, ice bath, centrifugation and removal of the supernatant. Resuspend and precipitate the 100 μL MNase Digestions Buffer. Add 0.25 μL Micrococcal Nuclease, mix it well for water bath. Add 10 μL of MNase Stop Solution to stop the reaction, followed by ice bath, centrifugation, and removal of the supernatant. 50 μL Lysis Buffer 2 was resuspended and precipitated, followed by ice bath, vortex, centrifugation and supernatant collection (H3K9 acetylation expression of the target protein in the sample was detected by WB method.)

5 μL of the supernatant obtained from the above steps was stored at −20°C. This is Input. 45 μL of the supernatant obtained from the above steps was put into a centrifuge tube containing IP Dilution Buffer. Positive control IP 10 μL Anti-RNA polymerase □antibody, Negative control IP 2 μL Normal Rabbit Ig G, Target-specific IP 5 μg antibody. For each IP, add 500 μL Diluted lysate to the plug spin column, add the primary antibody and let it stay at 4°C overnight. Add 20 μL of ChIP Grade Protein A/G plus Agarose to each IP, incubate for 1 h on a shaker, and discard the supernatant by centrifugation. Add 500 μL IP Wash Buffer 1 and discard the supernatant by centrifugation. Add 500 μL IP Wash Buffer 2 and discard the supernatant by centrifugation. Add 500 μL IP Wash Buffer 3 and discard the supernatant by centrifugation. Put the plug column back into the 1.5 mL centrifuge tube, add 150 μL IP Elution Buffer to wash resin, and incubate the resuspended magnetic beads. Add 6 μL 5M NaCl and 2 μL 20 mg/mL proteinase K by centrifugation and mix it well. Unfreeze Input, add 150 μL IP Elution Buffer, add 6 μL 5M NaCl and 2 μL 20 mg/mL proteinase K.

Add 750 μL DNA Binding Buffer to a DNA Clean-Up column, followed by centrifugation to discard the supernatant. Add the remaining sample to the same DNA Clean-Up column and add 750 μL DNA Column Wash Buffer. Centrifuge to discard the supernatant and centrifuge again. Add 50 μL DNA ElutionSolution, collect the purified DNA by centrifugation, and perform qPCR detection. The primer information is shown below. Translate system 20 μL, including forward primer, reverse primer, GoTaq®qPCR Master Mix, GoTaq®qPCR Master Mix. The amplification system was pre-denatured at 95°C for 10 min, denatured at 95°C for 15 sec, extended at 60°C for 1 min, and repeated for 50 cycles of reaction. The Ct data was obtained from the Bio-Rad PRISM Sequence Detection software in the Fluorescence quantitative PCR instrument.

### 2.6 Statistical analysis methods

The data was analyzed by SPSS20.0 software. The normally-distributed measurement results were expressed as mean ± standard deviation. The comparison between the two groups was performed by t test. The comparison between more than the two groups was analyzed by one-way analysis of variance. P<0.05 indicates statistically significant significance.

## 3. Results

### 3.1 Clinical sample information

Laboratory serological information such as age, transaminase (ALT), total bilirubin (TBIL), prothrombin activity (PTA) and HBV viral load of the enrolled subjects was counted (Table 1). The results showed that the three groups of samples were comparable in terms of gender, age, and BMI (P> 0.05), and HBV viral load in the treatment group decreased significantly (21).

**Table 1.** The clinical results between Control, HBV and Ta1-ETV treatment group (x±s).

| Indicators | Control | CHB | Ta1-ETV | P Value |
| --- | --- | --- | --- | --- |
| Patients (n) | 15 | 28 | 29 | / |
| Age<sup>a</sup> (y) | 35.34±7.65 | 39.71±13.20 | 47.71±13.20 | 0.211 |
| Gender (M/F) | 12/3 | 19/9 | 21/8 | 0.794 |
| BMI ( kg/m ) | 24.16±5.33 | 22.19±7.61 | 25.29±5.66 | 0.992 |
| HBsAg (+) | 0 | 28 | 29 | 0.151 |
| HBeAg (+) | 0 | 16 | 9 | 0.093 |
| ALT<sup>a</sup> (IU/L) | 23.6±2.9 | 986.5±82.7 | 42.3±6.6 | 0.000 |
| TBIL<sup>a</sup> (μmol/L) | 10.9±2.1 | 197.1±29.3 | 25.2±3.9 | 0.000 |
| PTA<sup>a</sup> (%) | 86.1±9.2 | 49.9±5.8 | 81.2±7.7 | 0.019 |
| HBV DNA<sup>b</sup> | ND | 7.71±1.1 | 0 | 0.032 |

ALT, alanine aminotransferase; TBIL, total bilirubin; PTA, prothrombin activity.
<sup>a</sup> For age and ALT, TBIL, PTA and HBV DNA (log<sub>10</sub> IU/ml), mean SD is shown for each group.
<sup>b</sup>Log<sub>10</sub> IU/mL

### 3.2 Correlation between anti-HBV therapy based on thymosin a1 and entecavir and TLR9 gene expression

In the previous work, targeting at the differential expression of Tolls in PBMCs of HBV-infected patients, we first examined the relationship between TLR9 gene expression and HBV DNA load by real-time quantitative PCR. The results are shown in Fig. 1A. There was a significant positive correlation between the expression of serum PBMCs TLR9 mRNA and the serum HBV DNA load in patients with chronic hepatitis B (n = 45). To investigate the effect of TLR9 on the pathogenesis of hepatitis B, we first explored the relative expression of TLR9 mRNA in healthy volunteers, HBV-infected patients and people treated with entecavir and thymosin a1. The results are shown in Fig. 1B.

**Fig. 1.**
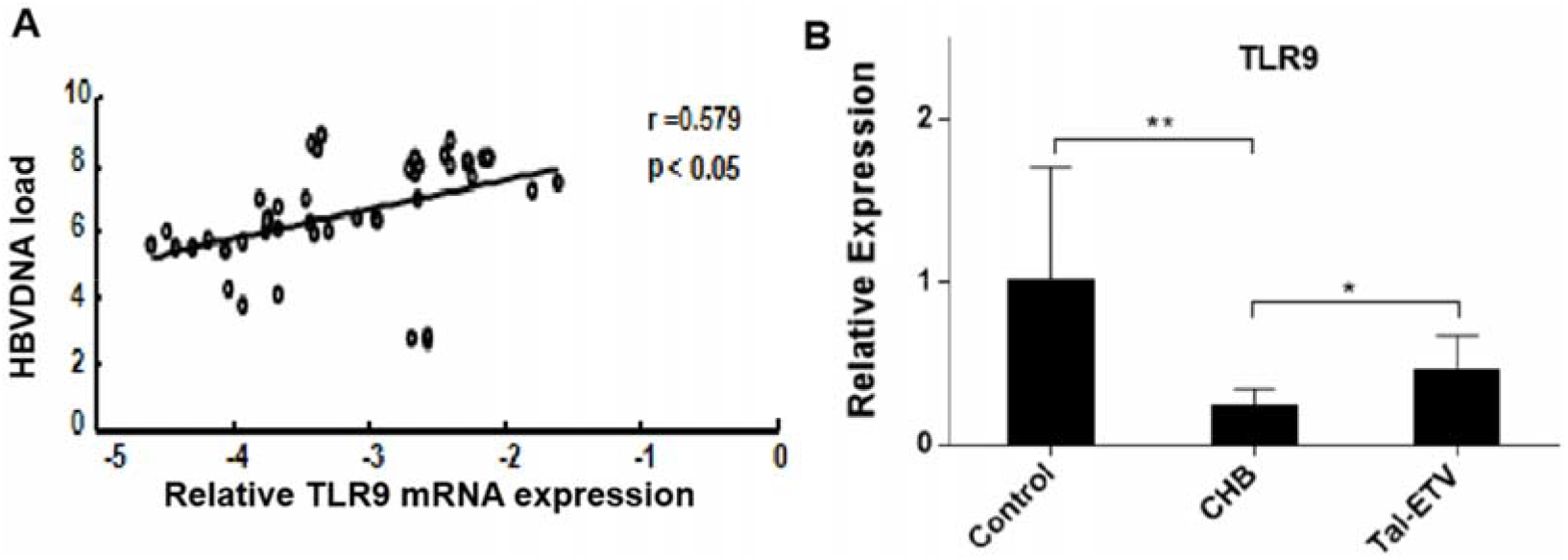
Correlation between expression of TLR9 mRNA and serum HBV viral load in patients with chronic hepatitis B (A), relative expression levels of TLR9 mRNA in different populations.

To this end, combining Elisa technology, we examined the expression of TLR9 protein at different treatment times, and the results are shown in Fig. 2. The results showed that compared with healthy people, patients with chronic hepatitis B had decreased serum TLR9 protein expression under HBV virus stimulation, but TLR9 protein expression increased after a certain period of treatment. This shows that TLR9 plays an important role in the treatment of chronic-on-chronic hepatitis B with entecavir and thymosin a1.

**Fig. 2.**
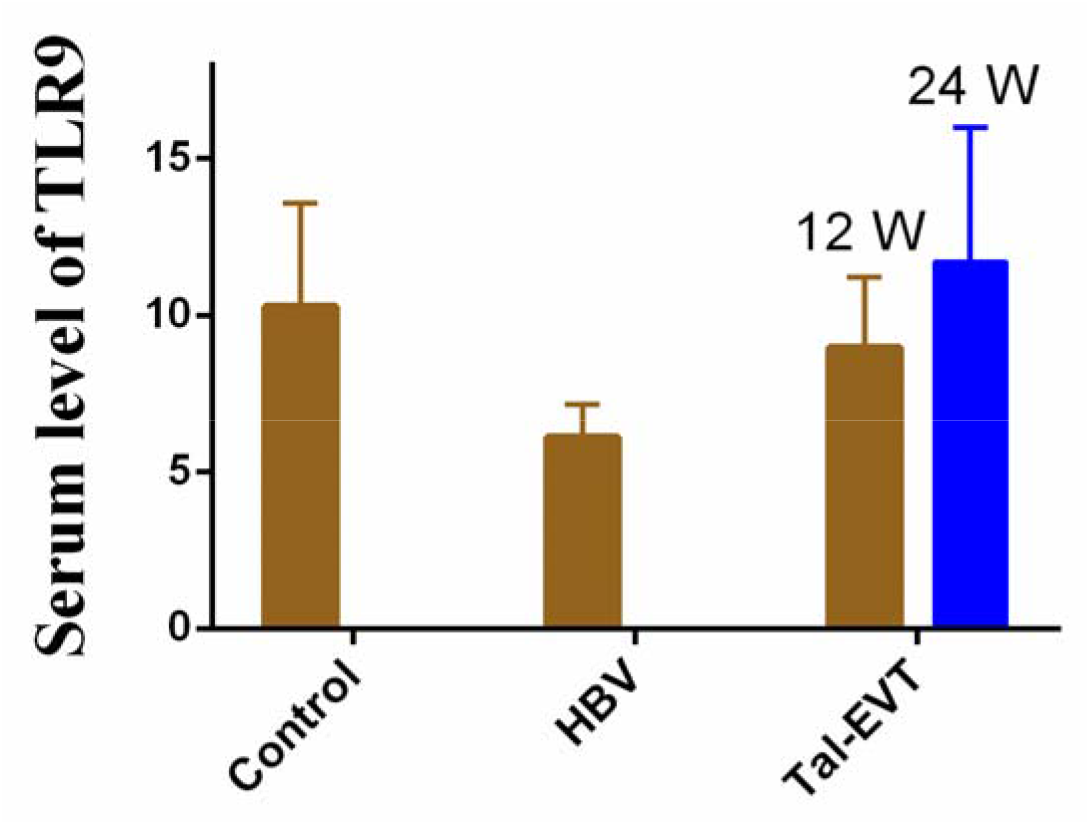
Comparison of serum TLR9 protein expression levels in different populations.

### 3.3 H3K9 acetylation modification of TLR9 gene during anti-HBV treatment with thymosin a1 and entecavir

Further, using ChIP experiment, we investigated the binding of protein H3K9Ac to the promoter region of TRL9 gene in the serum PBMCs of patients during the combined use of entecavir and thymosin a1. Table 2 shows the three predicted potential binding sites of TLR9 gene and protein H3K9Ac.

**Table 2.**
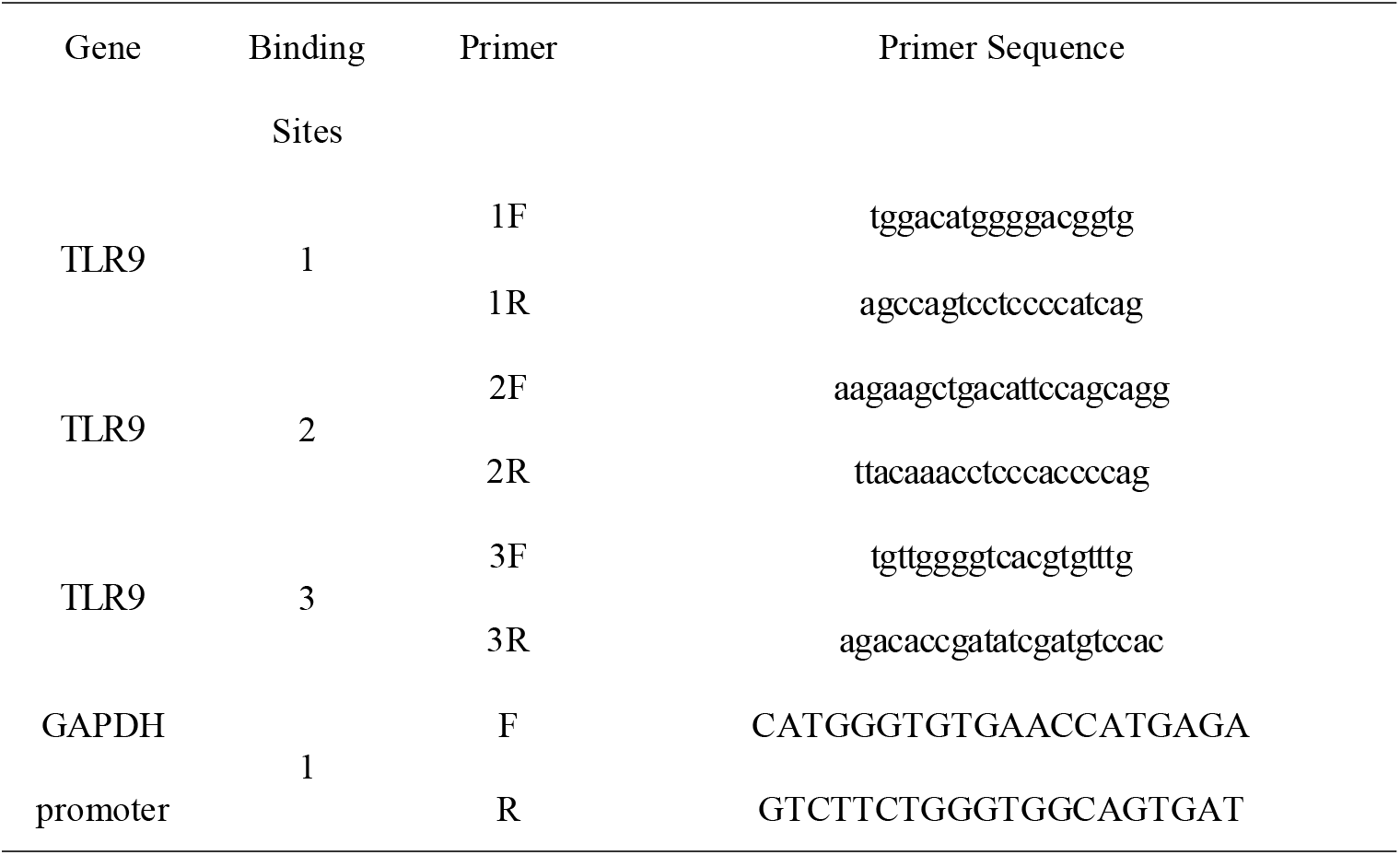
The gene-specific primers of the binding sites for H3K9Ac and TRL9.

The protein of the positive control was RNA polymerase II, and the gene sequence detected by qPCR was gapdhpromoter region. The protein of negative control was Rabbit IgG, and the gene sequence detected by qPCR was gapdhpromoter region and target gene indicators. As shown in Fig. 3, TLR9 gene has binding with H3K9Ac in serum PBMCc of healthy people, HBV-infected people and chronic hepatitis B patients treated with entecavir and thymosin a1. The data tested in each sample revealed that changes in enrichment rate of H3K9Ac and ChIP was disease-specific. It was found from the detection of 3 sites that differences in sites 2 and 3 were obvious between patients and normal people. DNA sequence analysis also revealed that H3K9Ac had the same changes in regional modification model in other cell models. This view is consistent with the finding that changes in histone modifications in fixed DNA sites induces gene regulation model.

**Fig. 3.**
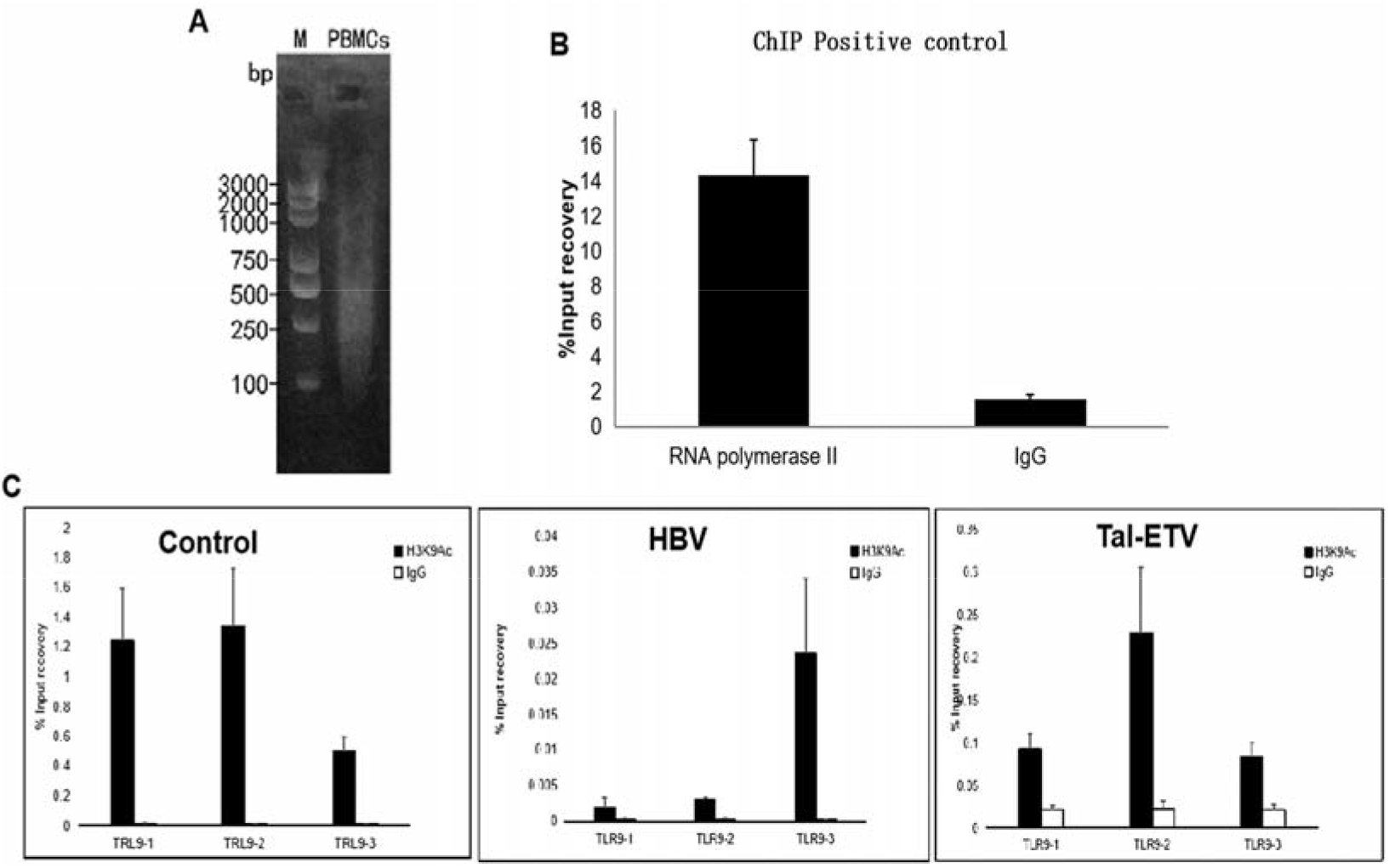
ChiP test to detect the binding of TLR9 gene to protein H3K9Ac in serum PBMCc of different populations. A) Fracture map of PBMCc cell chromatin extraction; B) Positive control of ChiP experiment; C) Binding status of TLR9 gene and protein H3K9Ac in PBMCc in different populations.

### 3.4 Detection of inflammatory factors during anti-HBV treatment with thymosin a1 combined with entecavir

To investigate whether H3K9 acetylation modification of TLR9 gene was a result of immune regulation activation during the virus clearance process of the combination therapy, we first tested expression levels of IL-6, IL-12, IFN-γ, and TNF-α in the serum of healthy volunteers, hepatitis B patients and patients with combination therapy, with results shown in Fig. 4.

**Figure 4.**
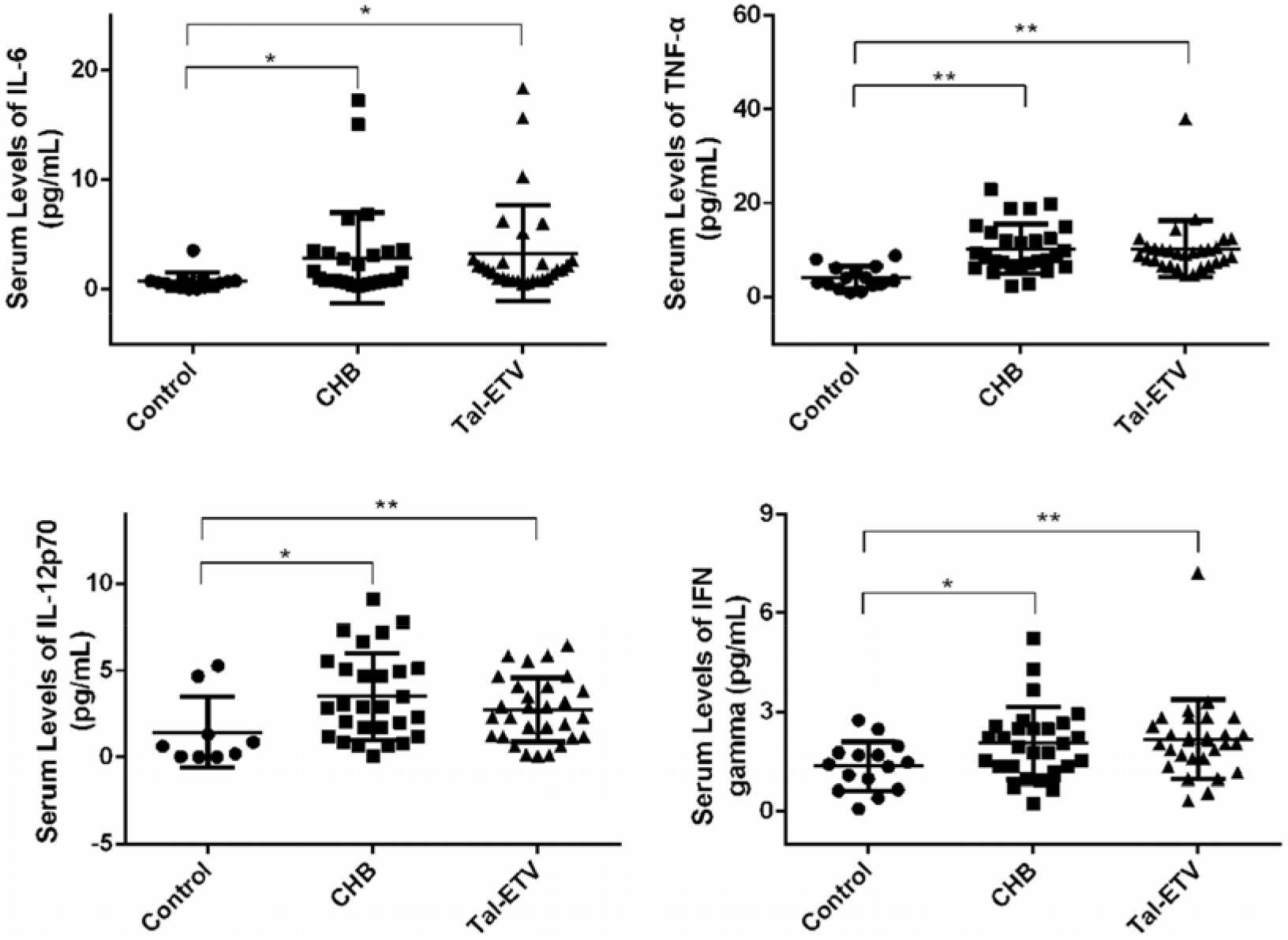
Mean (SD) serum levels of interleukin (IL)–6, IL-8, tumor necrosis factor (TNF)–r, and transforming growth factor (TGF)–ß in patients with chronic hepatitis B (CHB) infection, Tal-ETV treatment and healthy control individuals.

In order to explore the changes in immune balance during the treatment of chronic hepatitis B patients with entecavir and thymosin a1, we further examined the immune factors such as IL-6, IL-12, IFN-γ, and TNF-α at 12th, 24th and 48th week’s treatment cycles, with results shown in Fig. 5.

**Figure 5.**
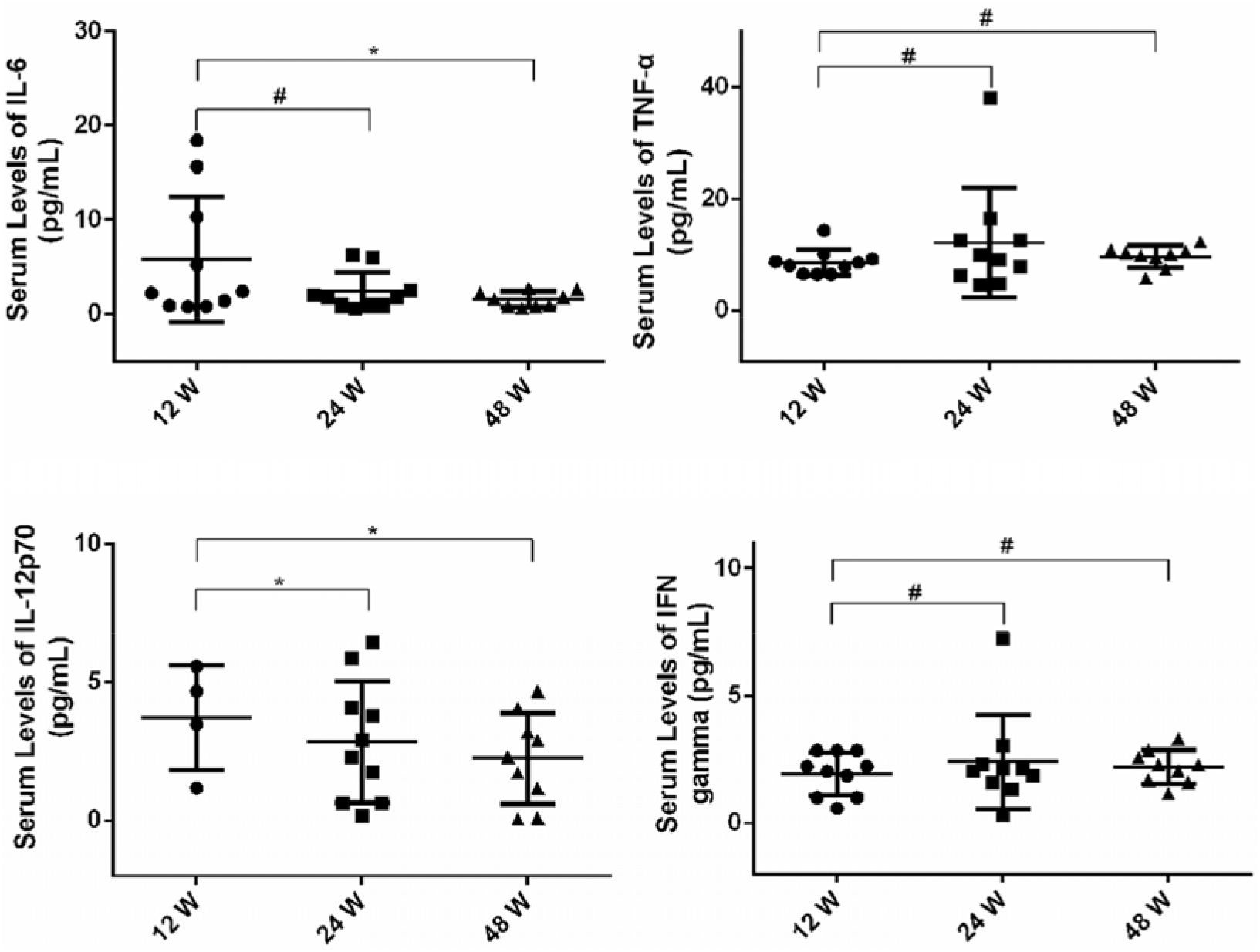
Mean (SD) serum levels of interleukin (IL)–6, IL-8, tumor necrosis factor (TNF)–i, and transforming growth factor (TGF)–ß in patients with different time after Tal-ETV treatment.

## 4. Discussion

At present, treatment of hepatitis B mainly adopts antiviral drugs to improve inflammatory response and viral infection of liver tissues, thereby improving clinical symptoms (4-5). However, there has been no perfect method for antiviral treatment of hepatitis B. Problems such as poor curative effect, high recurrence rate, and long treatment course still plague front-line clinicians. To this end, it carries extraordinary positive significance for us to explore new treatment methods for hepatitis B patients, clarify its mechanism of action, thereby improving the quality of life of patients with hepatitis B, improving quality of medical care and expanding the scope of medical services. Entecavir belongs to 2-deoxyguanine nucleoside analogue, which is a clinically approved drug by the US FDA for the prevention and treatment of hepatitis B. It is mainly used for active replication, liver histology or serologically verified treatments. Thymosin a1 is a polypeptide substance in the main active ingredient of thymosin, which can be used to improve cellular immune function. Thymosin a1 can help strengthen the body’s anti-infective ability, thereby reducing the immune damage to liver cells, achieving repair effects, inhibiting liver cell apoptosis, and protecting the liver. Clinical studies have found that treatment of patients with chronic hepatitis B by entecavir and thymosin a1 has significantly better efficacy in overall than entecavir alone, indicating that strengthening immune regulation during antiviral therapy can further strengthen hepatitis B virus clearance effect, and better improve patients’ clinical symptoms, enhance body immunity and curative effect. For this reason, exploring the molecular mechanism of entecavir and thymosin a1 in the treatment of chronic hepatitis B will provide an important theoretical reference for the proposal of more treatment schemes.

A large number of literature reports have found that TLR9 is activated in the immune cells of the body after stimulation by various viral infections and exerts its immune function by stimulating the immune system (18-19). TLR9 has the ability to identify HBV virus and initiate immune response. TLR9 expressed in cells is mainly located in the endoplasmic reticulum. Once infected viruses or bacteria are digested by intracellular lysosomes and endosomes, single-stranded DNA containing unmethylated CpG motifs will be released, which can recruit TLR9 for recognition and binding, thereby initiating the downstream immune activation response, leading to expression of multiple inflammatory cytokines and mediating immune response. HBV’s genomic DNA is a circular partial double helix structure with a length of about 3200 bp. Methylation-specific software was adopted to find nucleotide sequence containing CpG from the full-length HBV genome downloaded by NCBI. It was found that both HBV genome B genotype and C genotype contained a large number of CpG sites, and a CpG island was found, that is, the region contained CpG sites higher than the general frequency. During the replication or degradation of HBV virus, the presence of unmethylated CpG sequences in the open genome provides a theoretical basis for the recognition of TLR9 and initiation of immune responses. At present, there are also reports that TLR9 expression is inhibited in the infection process of chronic hepatitis B patients, causing immune escape.

In previous work, in response to the differential expression of Tolls on PBMCs in HBV-infected patients, we found that the expression of PBMCs TLR9 mRNA in serum of patients with chronic hepatitis B had a significant correlation with serum HBV DNA load (Fig. 1A). To investigate the effect of TLR9 on the pathogenesis of hepatitis B, we further explored the relative expression of TLR9 mRNA in healthy volunteers, HBV-infected patients and people treated with entecavir and thymosin a1. The results are shown in Fig. 1B. Taking the expression level of healthy volunteers as a control, TLR9 expression was significantly reduced in HBV-infected people, which was consistent with previous reports. HBV can down-regulate the expression of TLR9 mRNA and protein to reduce the production of immune factors. However, the relative expression of TLR9 mRNA increased significantly after combination therapy, indicating that epigenetic changes may occur in HBV infection and TLR9treatment. To this end, in combination with Elisa technology, we examined the expression of TLR9 protein at different treatment times, with results shown in Fig. 2. The results showed that compared with healthy people, patients with chronic hepatitis B had decreased expression of TLR9 protein in the serum under HBV virus stimulation, but TLR9 protein expression increased after a certain period of treatment. This indicates that TLR9 plays an important role in chronic-on-chronic hepatitis B treatment with entecavir and thymosin a1 (23).

In the treatment of chronic-on-chronic hepatitis B with entecavir and thymosin a1, the rise of TLR9 mRNA levels may be achieved through two pathways: mRNA transcription pathway and histone modification pathway. In the previous stage, the group compared acetylation modification state of histone H3K9 in peripheral blood CD4^+^T cells in whole genome promoter region under different disease states of chronic hepatitis B, finding that H3K9 acetylation modification regulation existed in 2 sequence regions of TLR9 (Ch3:52234647-52235096 and Chr3:52234897-52235346), suggesting that H3K9 acetylation modification of TLR9 might also play an important role in the occurrence and development of chronic hepatitis B. To this end, through ChIP experiments, we further investigated the binding of protein H3K9Ac to TRL9 gene promoter region in the patients’ serum PBMCs during the combination therapy of entecavir and thymosin a1 (27). As shown in Fig. 3, the data tested in each sample found that changes in the enrichment rate of H3K9Ac, ChIP was disease-specific and significant. From the detection of 3 loci, it was found that loci 2 and 3 had significant differences between patients and normal people. DNA sequence analysis also revealed that H3K9Ac has the same changes in regional modification model in other cell models. This view is consistent with the finding that changes in histone modifications in fixed DNA sites induce gene regulation model.

The pathogenesis of hepatitis B concerns the immune response and immune regulation disorders caused by hepatitis B virus infection of liver cells. A large amount of interleukin and tumor necrosis factor α can appear in the serum of patients with hepatitis B. Interleukins can promote the differentiation and proliferation of T cells, B cells, NK cells lymphokines and other activated killer cells, thus forming a dynamic immune regulatory network to maintain the body’s normal immune regulatory functions (32). The imbalance of this network has a close relationship with the occurrence and development of various diseases. With the help of liquid chip detection, the expression levels of IL-6, IL-12, IFN-γ and TNF-α in the serum of healthy volunteers, hepatitis B patients and patients with combination therapy can be obtained as shown in Fig. 4. Compared with healthy people, expressions of the four indicators after HBV infection are significantly increased, but decreases slightly after the combination therapy. In different treatment cycles, expression of IL-6 and IL-12 showed a significant downward trend; while expression of IFN-γ and TNF-α showed a slight upward trend, but P-value value was insignificant. The reason may be due to the small sample scale and too many confounding factors in peripheral blood samples.

To conclude, the results of this study show that H3K9 acetylation modification of TLR9 exists and plays an important role in patients with chronic hepatitis B (21). During the combination therapy with entecavir and thymosin a1, histone acetylation modification of TLR9 was significantly improved, which increased the expression of TLR9 at the mRNA and protein levels, and further regulated IL-6, IL-12 and other cytokines. The expression results are correlated with antiviral efficacy. The results confirm that TLR9 participates in the immune response of anti-HBV process using Ta1 combined with entecavir, which provides a new basis for the antiviral immune control theory of chronic hepatitis B, and has broad social needs and important clinical application significance.

## Acknowledgements

This work was supported by the National Natural Science Foundation of China (No. 81700528), Social Science and Technology Foundation of Dongguan (201750715001440).

## Authors’ contributions

Wang Ke, Zhong QY, Yin SC, Zhong JB, Du W and Li FW performed the experiments. Wang Ke and Zhu HP designed and directed the experiments. Zhu HP, Wang Ke and Cao HH wrote the manuscript.

## Ethics approval and consent to participate

All healthy human blood experiments were approved and conducted according to the experimental protocol authorized by the Ethical Committee of the Dongguan People’s Hospital.

## Patient consent for publication

Not applicable.

